# The virome of bats inhabiting Brazilian biomes: knowledge gaps and biases towards zoonotic viruses

**DOI:** 10.1101/2022.10.11.511773

**Authors:** Gabriel Luz Wallau, Eder Barbier, Alexandru Tomazatos, Jonas Schmidt-Chanasit, Enrico Bernard

## Abstract

Bats are hosts of a large variety of viruses including some that may infect other vertebrates and humans. Research on bat-borne viruses attracted significant attention in recent years mainly due to epizootics caused by viruses having bats as hosts. The characterization of the viral communities of bats was then prioritized, but despite increasing efforts, there are large disparities in the geographical ranges covered and the methodologies employed around the world. As a result, large gaps remain in our current understanding of bat viromes and their role in disease emergence. This is particularly true for megadiverse regions in Latin America. This review aims to summarize the current understanding about bat-viruses that inhabit Brazilian biomes, one of the most bat species-rich and diverse regions of the globe. Taking into account all known bat-associated viral families studied in Brazilian biomes, we found that almost half of all bat species (86/181 species) were not investigated for viruses at all. Moreover, only a small fraction of viral lineages or families have been studied more in depth, usually employing targeted methods with limited power to characterize a broad virus diversity. Additionally, these studies relied on limited spatio-temporal sampling and small sample sizes. Therefore, our current understanding of bat viral communities in the Brazilian biomes is limited and biased at different levels, limiting zoonotic risk assessments of bat-borne viruses. Considering these limitations, we propose strategies to bridge the existing gaps in the near future.

## Introduction

Bats (order *Chiroptera*) compose the second most diverse mammalian order with 1456 known species from 21 families (https://batnames.org) which participate in diverse ecological processes such as plant pollination, seed dispersion, soil renewal up to habitat modifications(1). Bats are also reservoirs of a large diversity of zoonotic and non zoonotic viruses (2–4). The identification of specific bat species as reservoirs of human pathogenic viruses (Nipah paramyxoviruses and SARS-coronaviruses) propelled increasing efforts to characterize zoonotic viruses with epidemic potential that infect and are transmitted by bats(2). Many studies underscored that Chiropterans are more prone to carry and transmit zoonotic virus to humans(5–8). On the other hand, non-bat intermediate amplifying hosts and the characterization of zoonotic viruses hosted by other mammalian species suggest that a complex transmission chain in different animal hosts ultimately leads to spillover to humans(2, 9, 10). Where there is evidence of direct transmission of bat viruses to humans, human activities had a prominent role promoting contact with bats(11–15). A recent study showed that the number of zoonotic viruses found in mammalian orders is proportional to the number of species of each respective order, suggesting that bats are not special pathogen reservoirs and that they host a large diversity of viruses because of its high species diversity(16). There is still no consensus if bats do carry and spread more human-pathogenic viruses than other mammalian orders(17).

Definitive answers to the role of bats as reservoirs of zoonotic viruses - and its associated risks - ultimately lies on the full understanding of bat’s viromes, ecology and their interaction with other vertebrates. Several studies sampled wild bats and populations to more broadly characterize its virus diversity. However, there are large disparities in study efforts in different regions of the world, since the vast majority of virus studies were largely performed in Asia, Africa and Europe, with a lower number of studies in Oceania and North America(3, 18, 19). Moreover, Central and South America, among the most bat species-rich regions of the globe, are comparatively poorly studied (20). Another hardly addressed issue in viral studies on bats is the limited representative sampling, especially considering the large habitat range and population size variation of bats and the infection rate fluctuations through time(21). Therefore, our understanding of bats viromes is restricted due to biases at different levels..

Bats virome characterization is also biased when it comes to molecular methodologies employed for viral detection, once the majority of virus detection methods were developed to target single viral lineage of zoonotic concern and not to capture the full diversity of bat-viruses(20). These methodologies rely on previous information about the pathogen molecules (22) which contrasts with our scant knowledge of the animal’s virome(23). Nowadays, high throughput methodologies (hereafter HTS) is one of the most promising approaches for unbiased viral nucleic acid detection, allowing a broad description of viral genomes associated with any sample and requiring no previous knowledge about the infecting viral pathogen(24). HTS virome studies generated major breakthroughs, revealing thousands of new viral lineages and families(25), but also brought to light that the large majority of animal viruses still awaits to be formally characterized(26–28).

In this review we will address the current understanding of viruses detected in bats inhabiting Brazilian biomes (181 known species - https://www.sbeq.net/lista-de-especies). We highlight the currently limited and biased character of research of Brazilian bats virome. Additionally, we summarize the methodologies used for characterization of bat viruses, showing that most studies used targeted low throughput assays. Finally, we propose methodological approaches to address existing gaps.

## Material and Methods

### Database construction

We screened Pubmed, Google Scholar and Scopus scientific literature databases for manuscripts containing the terms “bat-borne viruses”, “viruses AND bats”, “viruses AND Chiroptera”, “virome”, “South American bats’’, “Brazilian bats”, “Rabies”, “Coronavirus”, “Hantavirus” until May 2022. All the search terms were known to be represented in the published literature of viruses detected in South American bats. Additionally, we added manuscripts in Portuguese and Spanish which could be missing in the searches based on English keywords. Although the review was focused on bats inhabiting the Brazilian biomes/territory, some species have larger distribution areas covering other South and Central American countries, and studies conducted other South American countries other then Brazil were also included. We built a reference library of 87 publications, for which 81 remained after filtering for bat sampling or full publication accessibility (**Supplementary File 1 and 2**).

For each study we extracted title, year of publication, viral family/lineage studied (specific viral families or several for virome studies), country, state, bat species, number of sampled individuals per species, number of positive individuals, methodology of virus detection, sampling strategies (longitudinal or single sampling) and type of sample (tissue or body fluid). We extracted data to species level for viruses and bats, and used family-level taxonomy for visualization. The methodologies employed for virus detection and characterization were classified according to throughput: low throughput (LT), high throughput (HT) and studies using both methodologies (LT + HT). We detected the following methodologies used in the published articles: Virus Isolation through Intracerebral Inoculation of Mice (VIIIM), the Virus Isolation in Cell Culture (VICC), Electron Microscopy and Antigenic Profile (ELANT), Western Blot (WB), Direct Immunofluorescence Antigen Monoclonal Antibodies (DIAMA), Enzyme-Linked Immunosorbent Assay (ELISA), Strain-specific PCR (SS-PCR), Family/Subfamily Degenerate PCR (FSD-PCR), Nested PCR (N-PCR), DNA Virome (DNAVIR), RNA Virome (RNAVIR) and Full Virome (FVIR) (**Table 1)**.

**Table 1.**
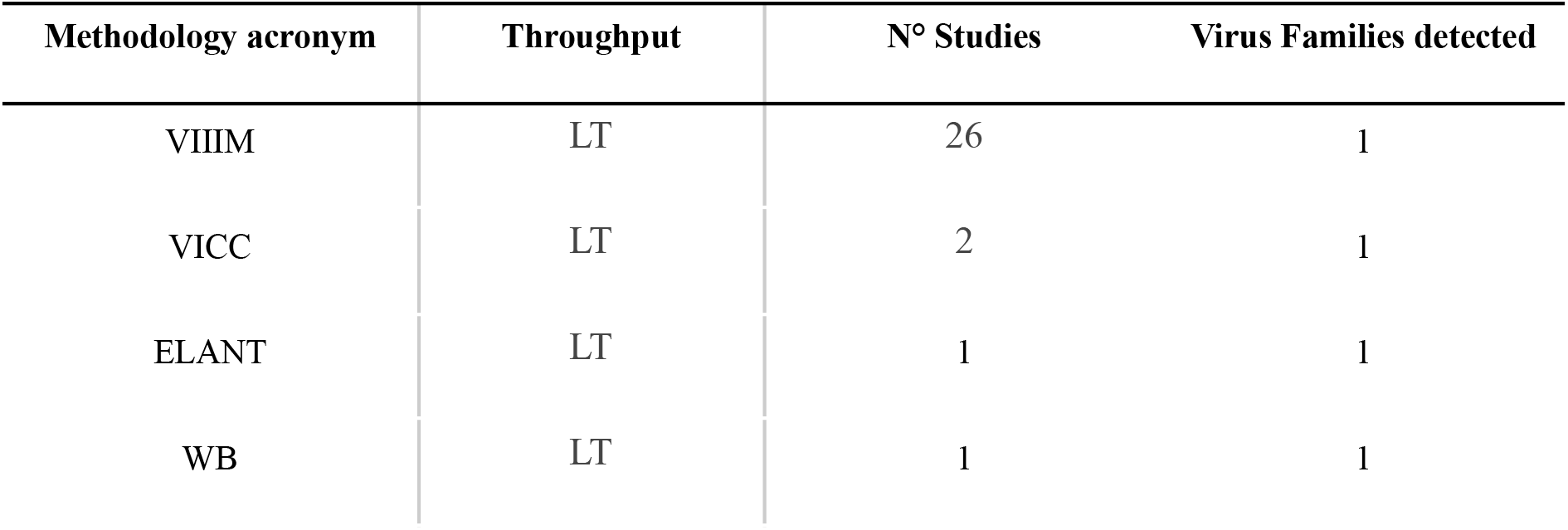
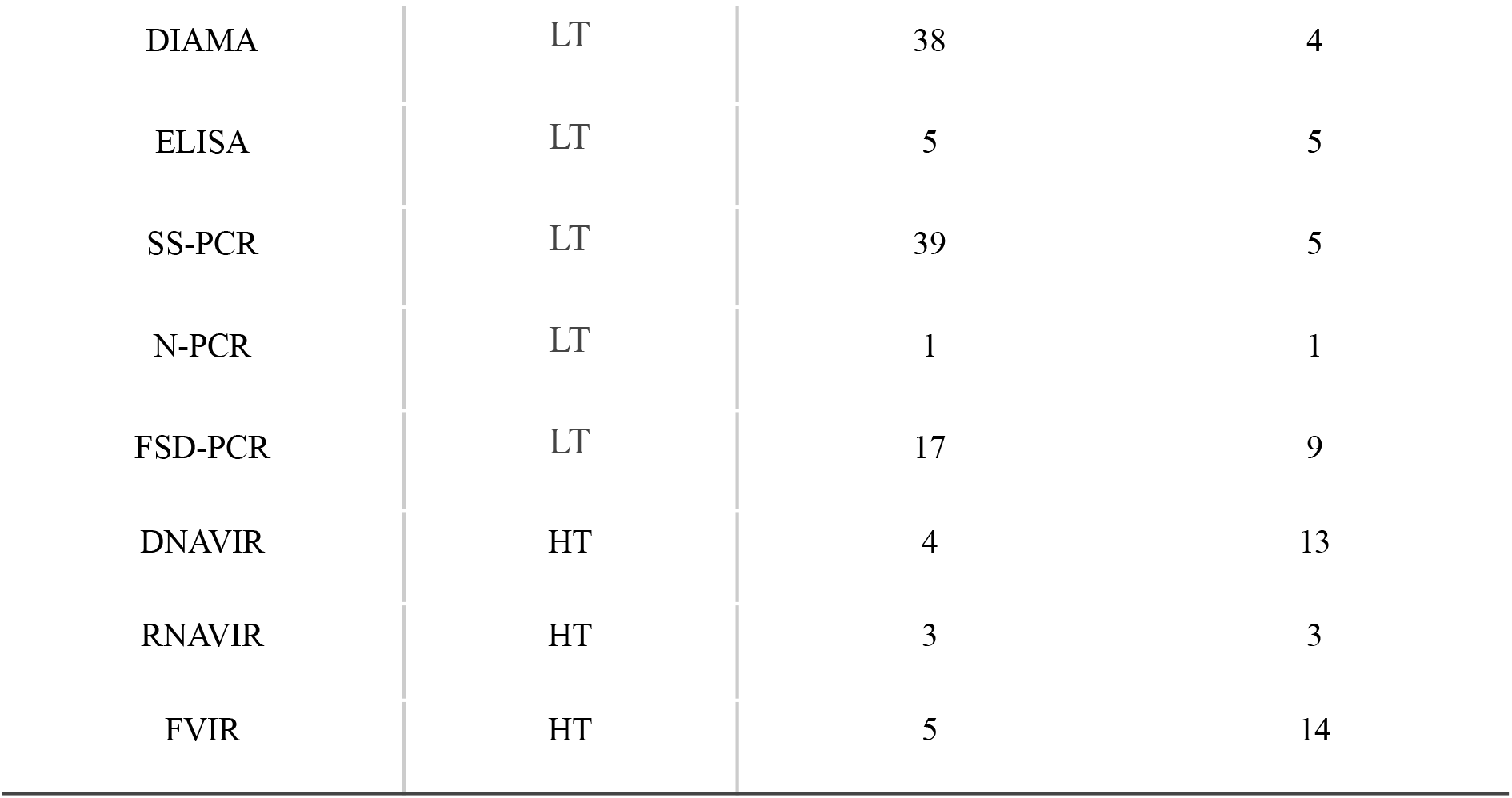
Methodologies employed in 81 studies with the number of studies using each methodology and the number of virus families identified. Methodologies are presented in a descending order from lower to higher throughput in terms of viral diversity characterization. Virus Isolation through Intracerebral Inoculation of Mice (VIIIM), the Virus Isolation in Cell Culture (VICC), Electron Microscopy and ANtigenic Profile (ELANT), Western Blot (WB), Direct Immunofluorescence Antigen Monoclonal Antibodies (DIAMA), Enzyme-Linked Immunosorbent Assay (ELISA), Strain-specific PCR (SS-PCR), Family/Subfamily Degenerate PCR (FSD-PCR), Nested PCR (N-PCR), DNA Virome (DNAVIR), RNA Virome (RNAVIR) and Full Virome (FVIR).

A number of studies have sampled and characterized viruses from bat excrements (guano) where many virus genomes were likely derived from food sources, internal or external bat microbiota. Therefore, we only extracted information of viruses that where known to infect vertebrates, irrespective of the original sample (guano, oral/anal fluids or tissues).

### Nucleotide database search and information extraction

In order to assess the available virus sequence information, we retrieved the number of entries per bat species and virus family from ZOVER database (http://www.mgc.ac.cn/cgi-bin/ZOVER/main.cgi - last accessed on May 2022)(29) using the most up-to-date taxonomy (https://www.sbeq.net/lista-de-especies). This database was chosen because it is the only database that compiled all available viral genetic information found in bats.

Figures were rendered with R statistical language (https://www.r-project.org/) employing packages such as: ggplot2(30), webr 0.1.2(31), dplyr 1.0.9, tidyr 1.2.0, ggpubr 0.4.0, reshape2 1.4.4, ggforce 0.3.3, ggridges 0.5.3, treemapify 2.5.5, ggdist 3.1.1, gghalves 0.1.3, packcircles 0.3.4, ggrepel 0.9.1, patchwork 1.1.1, ggvenn 0.1.9(32). Maps were rendered using QGIS version 3.26(33).

## Results and discussion

### Studies of bat viruses

Studies of bat viruses in Brazilian biomes were mostly performed from 1990 to the present (**Supplementary File 3**). From the 81 published studies analyzed, there is a clear focus on *Rhabdoviridae* and *Coronaviridae*, with 43 and 13 studies, respectively. The majority of studies focusing on *Rhabdoviridae*, and more specifically on the *Rabies* virus (RABV), were likely motivated by concerns about fatal spillover events to humans in Brazil once different bat species are described as important RABV reservoir(34–36). Few studies (1-3) have focused on *Orthomyxoviridae, Paramyxoviridae, Hantaviridae, Herpesviridae* and *Adenoviridae* (**Supplementary File 3**).

### Bats and their viruses in Brazilian biomes

Of the 181 bat species known to exist in Brazilian biomes, 95 were included in at least one study, while 86 species (47.5%) were not screened for viruses so far (**Figure 1A**). The sampling biases are also reflected at the family level: Phyllostomidae (the most diverse family in Brazil with 92 species is represented by 51 species, while 41 species (44.1%) were not studied so far (**Figure 1A**). Some less diverse bat families such as Thyropteridae (5 species) were not investigated at all (**Figure 1A**).

**Figure 1.**
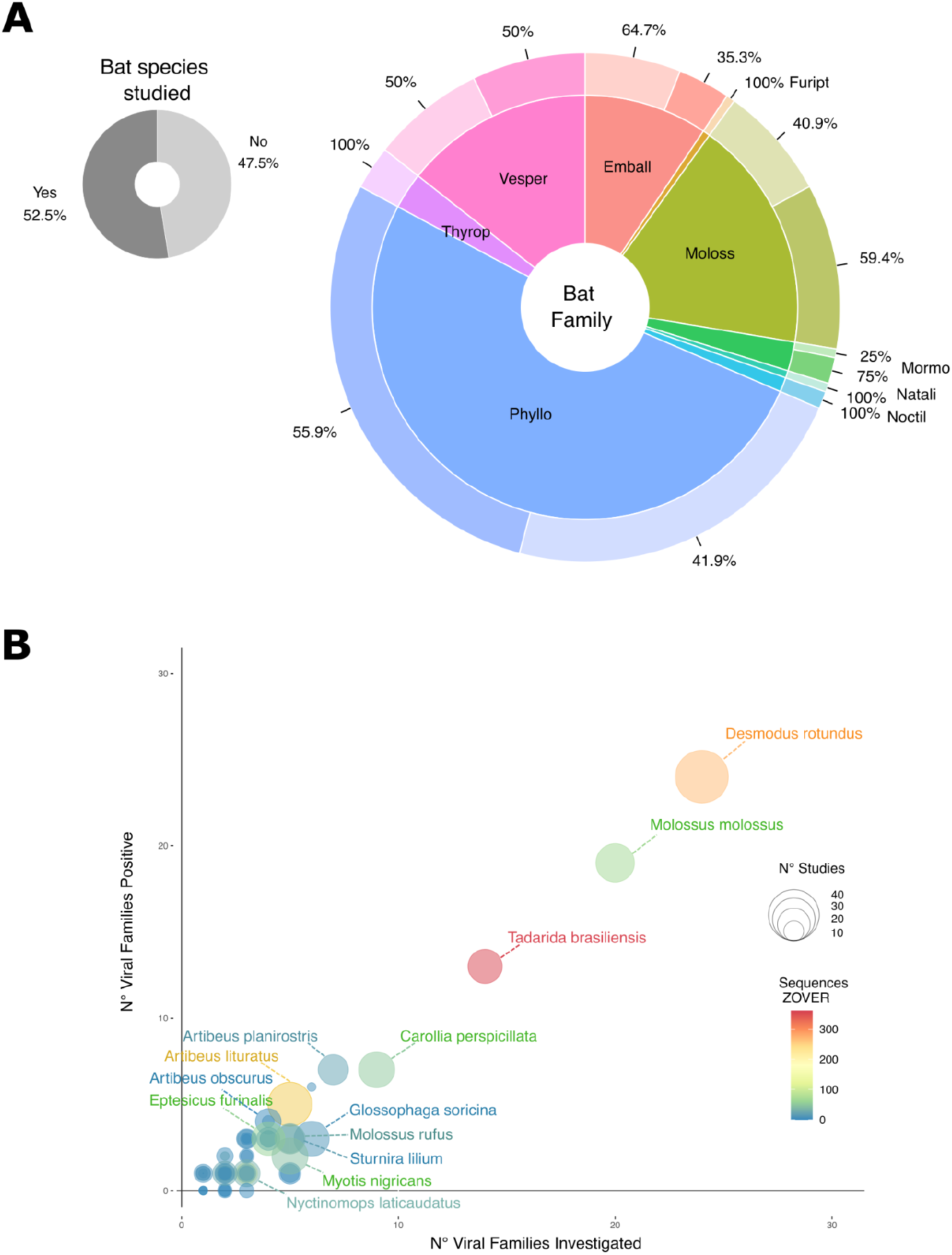
Amount of virus studies on bats of Brazilian biomes. **A** - Proportion of 181 known bat species from Brazilian biomes stratified per family that were included (dark gray) or not (light gray) in at least one study. Larger donut plot shows the proportion of studied species per bat family. Bat families are: Phyllo - Phyllostomidae, Moloss - Molossidae, Emball - Emballonuridae, Vesper - Vespertilionidae, Thyrop - Thyropteridae, Mormo - Mormoopidae, Natali - Natalidae, Noctil - Noctilionidae. **B -** Correlation between number of virus families detected or investigated and number of virus families found in the reviewed studies. Bubble size stands for the number of studies covering each bat species and bubble color represents the number of virus sequences of the corresponding species available in the ZOVER database.

Among the 95 species included in virus study, there is a clear over-representation of some important zoonotic virus reservoirs such as the vampire bat *Desmodus rotundus* (**Figure 1B**) which was screened for viruses in 44 of 81 studies (56.79%) reviewed (**Figure 1B**). This species is known to be the main sylvatic RABV reservoir in Brazil and the Americas (37), with infections of cows, dogs and humans traced to the *D. rotundus* RABV lineage (38, 39). In Brazil, rabies epidemiology shifted from a zoonotic infection mainly driven by domestic animals (dogs) to sporadic spillovers to cows and humans caused by lineages of bat origin, after effective prophylactic vaccination of rabid dogs were conducted country-wide (35, 38). Thirteen species were included in 10 or more studies (named species in **Figure 1B)**. All these species were detected as positive for different lineages of RABV (38, 40–46). Therefore, there is a visible sampling bias towards certain bat species. We found a correlation between the number of viral families investigated and number of viral families found for all species herein reviewed (**Figure 1D**), as well as between the number of virus families listed in this review and the sequences available from the ZOVER database (**Figure 1E**). Therefore, the larger the effort to sample a given species, the higher the likelihood of finding previously undetected virus families.

### Spatial sampling bias of bats in Brazil

The number of bats species known to occur in each Brazilian state ranges from 40 to 135 species (**Figure 2A and B**). However, bat sampling with the aim of studying viruses is highly heterogeneous. The number of bat species included in at least one study varied from 0 to 47 species. In 12 of 27 Brazilian states no species were analyzed (**Figure 2C**). Moreover, the ratio between the number of bat species studied and the total species inhabiting the respective states also differed. Only in São Paulo state more than 50% of the known bat species were assessed for virus infections (**Figure 2D**). For the remaining states, a maximum of 25% of known bat taxa were investigated (**Figure 2D**). From this we can observe that a spatial bias studies targeting the bats living predominantly in coastal states (**Figure 2C**).

**Figure 2.**
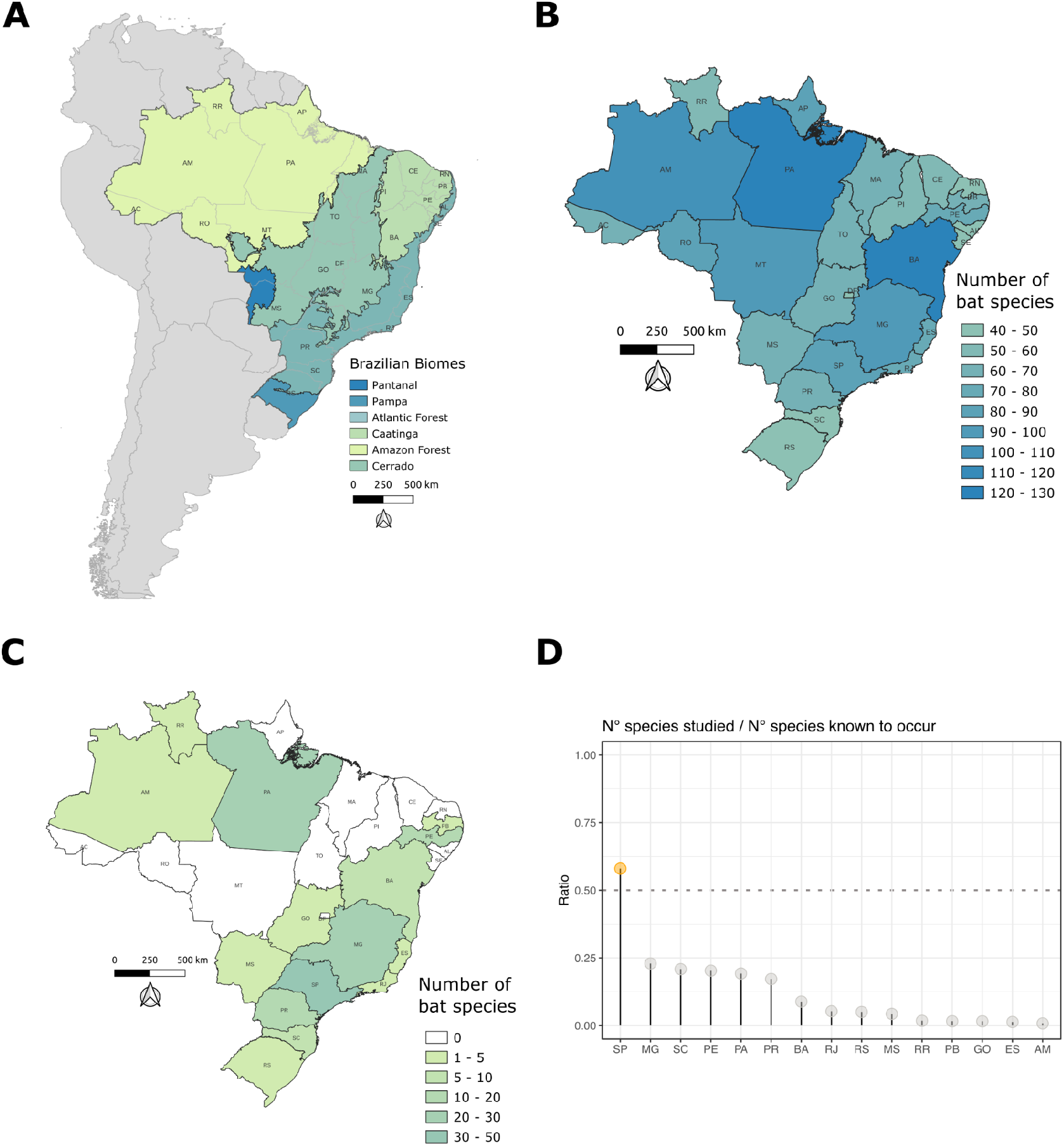
Brazilian biomes and bat species sampling effort on virus studies. **A -** Six Brazilian biomes covering different Brazilian states. **B -** Occurrence of known bat species per Brazilian state. **C -** Number of bat species investigated per state. **D -** Ratio of number of bat species investigated divided by the number of known species per state. Dotted line represents the ratio in which half of the species from a state are studied for viruses.

### Virus families studied

Altogether 17 RNA virus families and 14 DNA virus families were detected in our review and the ZOVER database (**Figure 3A and B**). Four families of DNA virus and four of RNA viruses were detected only in the review database, while two RNA virus families were found only in the ZOVER database. We have also found differences in the number and species recorded as positive for a viral infection/exposure. In the review compiled database we found 31 families in 62 of the 95 bat species studied representing 5 of 9 chiropteran families living in Brazil: Phyllostomidae (38), Molossidae (14), Vespertilionidae (9), Mormoopidae (2) and Emballonuridae (1). In the ZOVER database, evidence of infection was recorded for 22 virus families in 58 of 95 bat species. Here, the bats represent 6 of the 9 bat families recorded in Brazil: Phyllostomidae (24), Molossidae (15), Vespertilionidae (11), Emballonuridae (3), Mormoopidae (3) and Natalidae (1) (**Supplementary File 4**).

**Figure 3.**
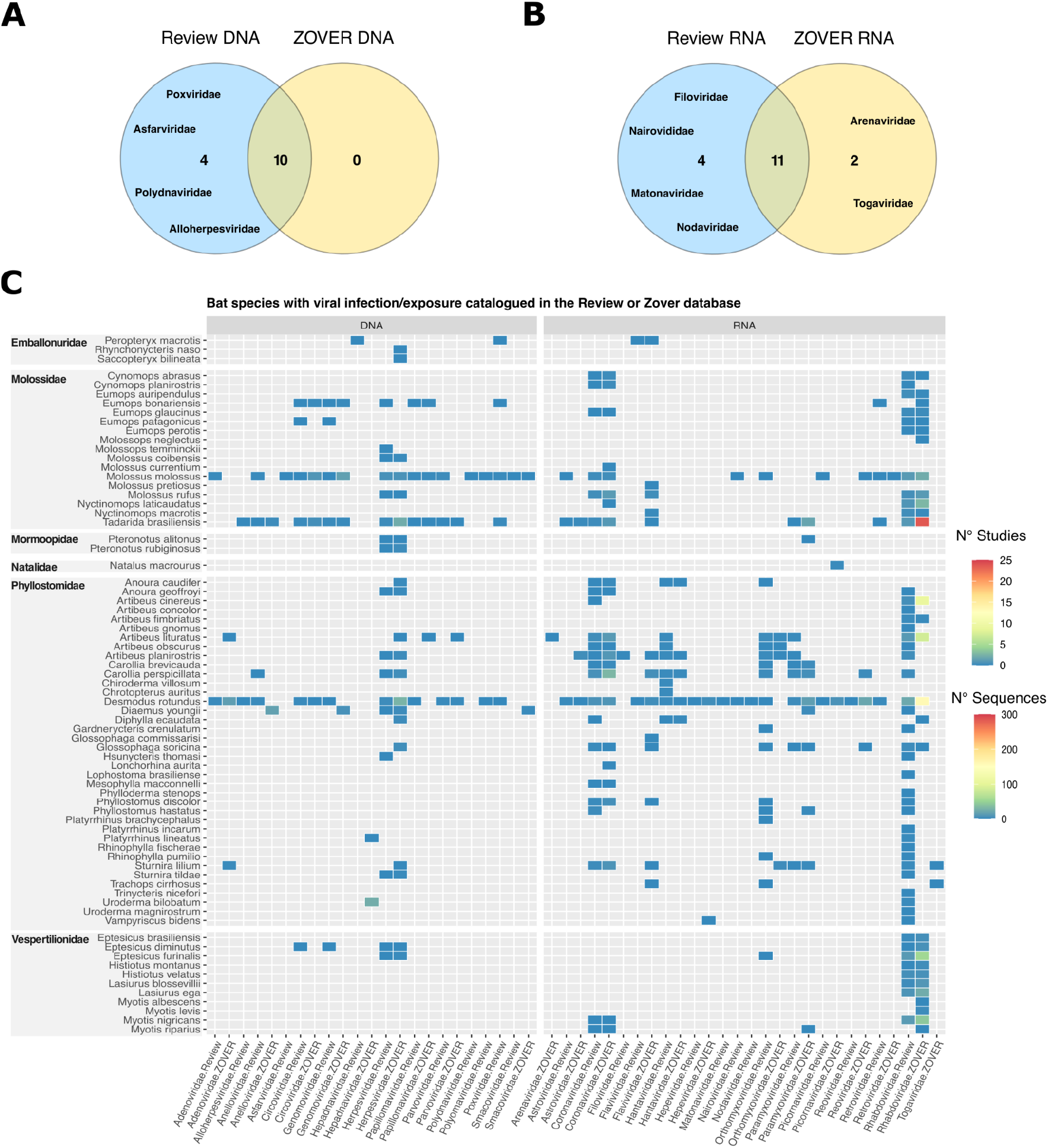
Bats inhabiting Brazilian biomes with evidence of viral infection/exposure obtained from studies cataloged in this review’s database and/or with sequences available in the ZOVER database. **A and B -** Venn diagram showing the number of virus families recovered in both datasets. **C -** Heatmap depicting the number of studies where evidence of viral infection is available and the number of virus sequences from each bat species per virus family (sorted by genome type).

We found a clear dominance of RNA virus data over DNA viruses available in both our review database and in the ZOVER database. The 5 most detected RNA virus families are *Rhabdoviridae* (47 bat species positive), *Coronaviridae* (21), *Orthomyxoviridae* (16), *Paramyxoviridae* (8) and *Hantaviridae* (9), while the remaining RNA virus families were detected in 5 or fewer bat species (**Figure 3C**). The 5 most covered virus families in the ZOVER database are: *Rhabdoviridae* (29), *Coronaviridae* (23), *Flaviviridae* (13) and *Paramyxoviridae* (10), while the remaining virus families were detected in fewer than four bat species. *Herpesviridae* was the only DNA virus family detected in more than 6 bat species with 17 species positive in the review database and 21 species positive in the ZOVER database (**Figure 3C**). The remaining DNA virus families were detected in 6 or fewer bat species.

These discrepancies in infected host diversity observed between our review database and ZOVER can be explained by two key differences. While our review cataloged studies using a large array of methodologies for viral detection, including the ones generating no genetic information (i.e. ELISA, DIAMA), the ZOVER database is based only on virus genetic information available from other public databases. This means that we were able to detect both infection (presence of RNA/DNA and antigens) and exposure (antibodies) of bat hosts in the review database, while the ZOVER database only provides information about the infection status of the host. Moreover, there are several examples of virus sequences available on the ZOVER database tagged as “*unpublished*” (257 entries, accession date July 1st, 2022), indicating that this data was publicly released with no peer review. Nevertheless, the majority of the bat species have not been comprehensively studied once most virus families were detected only in 1-3 bat species (**Supplementary File 5)**.

The total number of bat species sampled and studied is important information for evaluating virus family host range. However, only the host species that tested positive were reported in a number of manuscripts focused on RABV (**Supplementary File 2**) and such information can not be recovered from the ZOVER database. Therefore, we used the detection of virus families per host species as a proxy for the number of studied species from each bat family. This likely leads to an underestimate of the total host species/specimens investigated. Additionally, several virus families (12 of 29, DNA - *Smacoviridae, Anelloviridae, Papillomaviridae, Poxviridae, Alloherpesviridae, Asfarviridae, Polydnaviridae, Papillomaviridae* and *Parvoviridae*, and RNA viruses - *Retroviridae, Nairoviridae* and *Picornaviridae*) were only characterized using HTS methods, screening a limited number of individuals and species (see **Methodologies for virus detection**). Overall, these results highlight a very limited host range investigated for the majority of virus families studied.

### Infection status and temporal sampling strategies

The temporal sampling and the number of individuals sampled per species is heterogeneous. Longitudinal sampling was conducted only in 46 of 81 cataloged studies. Moreover, 27 of the 46 studies with longitudinal sampling have focused on RABV (**Supplementary File 6**). A recent meta-analysis of bat coronaviruses surveillance identified longitudinal repeated sampling (i.e. sampling the same site and bats populations multiple times) as the most significant determinant for viral detection (20), suggesting that the temporal variation in bat sampling is a key-component to take into consideration for viral surveillance. Regarding the infection status and prevalence we retrieved such information from 26 of the 81 studies only (**Supplementary File 6**). The lack of sample size information (total number of individuals collected) hindered estimates of infection prevalence for the large majority of species (**Supplementary File 6**). Virus infection status of each individual or population is highly variable depending on several factors such as the host immunological barrier acquired from previous infections, the number of susceptible newborns in a population and specific physiological, ecological and life history factors(15).

Implementing a sampling strategy that allows a broad assessment of bat-viruses is also challenging due to the diverse ecological patterns of viruses. Viruses interact with their hosts in various ways, e.g. pathogenic viruses cause acute and short-term infections, other lead to persistent/chronic infections. In each case, infectious status of individuals and populations may vary through time following virus-specific, tissue-specific replication and shedding(2). Hence, infection estimates may be affected by the sampling strategy (longitudinal vs cross-sectional). For instance, non-pathogenic viruses that develop a mutualistic interaction with their host may become highly prevalent in a population. In such cases, cross-sectional sampling schemes should still be effective to detect the virus, since a large fraction of individuals are infected at distinct timepoints. On the other hand, pathogenic viruses that induce acute infection and long lasting immunological response (weeks to months) usually increase in prevalence only during short periods of time. Therefore, such viruses can be more easily detected during ongoing sporadic outbreaks, when a large fraction of the host population is infected. The virus may go undetected if sampling does not take into account host physiological traits that influence prevalence in time (e.g. antibodies, starvation etc.). Coronaviruses,, for instance, show remarkable variance in fecal shedding, ranging from 0% to 80% in Old World bats(47). Paramyxoviruses such as Henipa viruses have been shown to be discharged in a pulse-like pattern, tightly linked to bat physiology status and waning of maternal antibodies in newborns(48). Filoviruses (e.g. *Marburg virus)*, also vary in prevalence, infection peaking during the birthing season of *Rousettus aegyptiacus*(49).

Different tissue tropism and route of viral transmission are additional factors impacting the sampling strategy. Collecting oral swabs provides information about viruses that are excreted in oral/nasal fluids but not about non excreted viruses or excreted through a different route, providing only a restricted view of the virome. Similarly, studies performing sampling of feces may characterize only viruses that are excreted through bats’ digestive tract along with a large diversity of viruses that are derived from food sources, but do not infect bats(50–52). The cataloged studies of bats from Brazilian biomes used various tissue types chosen with *a priori* knowledge of the target virus infectious dynamics and shedding. One illustrative example is RABV, investigated in 43 studies using euthanasia and collection of brain tissue, as the gold standard methodologies for RABV detection required numerous infectious particles for intracerebral inoculation and isolation (**Supplementary File 6**). Four of 13 studies investigating coronaviruses employed oral/nasal/anal swabbing due to the upper respiratory and intestinal replication and known viral sheeding route though oral/anal fluids (**Supplementary File 6**). It is important to note that three studies that screened coronaviruses in bats inhabiting Brazilian biomes used various tissues of internal organs from convenience samples obtained in RABV surveillance programs (**Supplementary File 6**). Convenience samples are known to be biased and generate misleading results for pathogen and disease prevalence (53).

Species populations and metapopulation size should be considered key driving factors in study design since virus prevalence is population size-dependent and host’s biology features can affect the likelihood of detecting positive individuals. However, sampling a wide range of bat sample types is only possible via euthanasia, which should be carefully considered in order to reduce the impact on the population and/or species. Some bat species have considerably low population size and sampling too many individuals may have a drastic impact on bat populations, leading to local extinction and likely long-term impact on these species.

Overall, studies on bats in Brazil are very heterogeneous in their temporal sampling strategy and the type of analyzed material. However, in order to broadly characterize bat viromes, flexible sampling strategies including systematic longitudinal sampling of representative individuals/populations and targeting different tissues must be adopted.

### Methodological approaches for virus characterization

Various methodologies were employed for characterization of viruses in Brazilian bats. The large majority of the studies employed low throughput methodologies (LT) (**Supplementary File 3 and Figure 4A**) such as VIIIM, that is based on the maceration of bat tissues and experimental intracerebral inoculation of mice(54). Positive VIIIM is followed by validation using direct immunofluorescence of RABV derived antigens with monoclonal antibodies (DIAMA - the most used methodology along with SS-PCR)(55) (**Table 1, Figure 4A**). Studies focused on *Coronaviriae* (11 studies) used only amplicon-based methodologies varying from strain-specific PCR (SS-PCR) and family/subfamily-level degenerated RT-PCR (FSD-PCR) (**Figure 4A**). While studies on other families employed a mix of different LT methods (**Figure 1A**). Twelve studies employed high throughput nucleic acid sequencing methodologies (FVIR, RNAVIR and DNAVIR) (**Figure 4A**). These studies employed complementary low throughput methodologies such as FSD-PCR, SS-PCR, ELISA, DIAMA and VICC (**Figure 4A** and **Supplementary File 6**).

**Figure 4.**
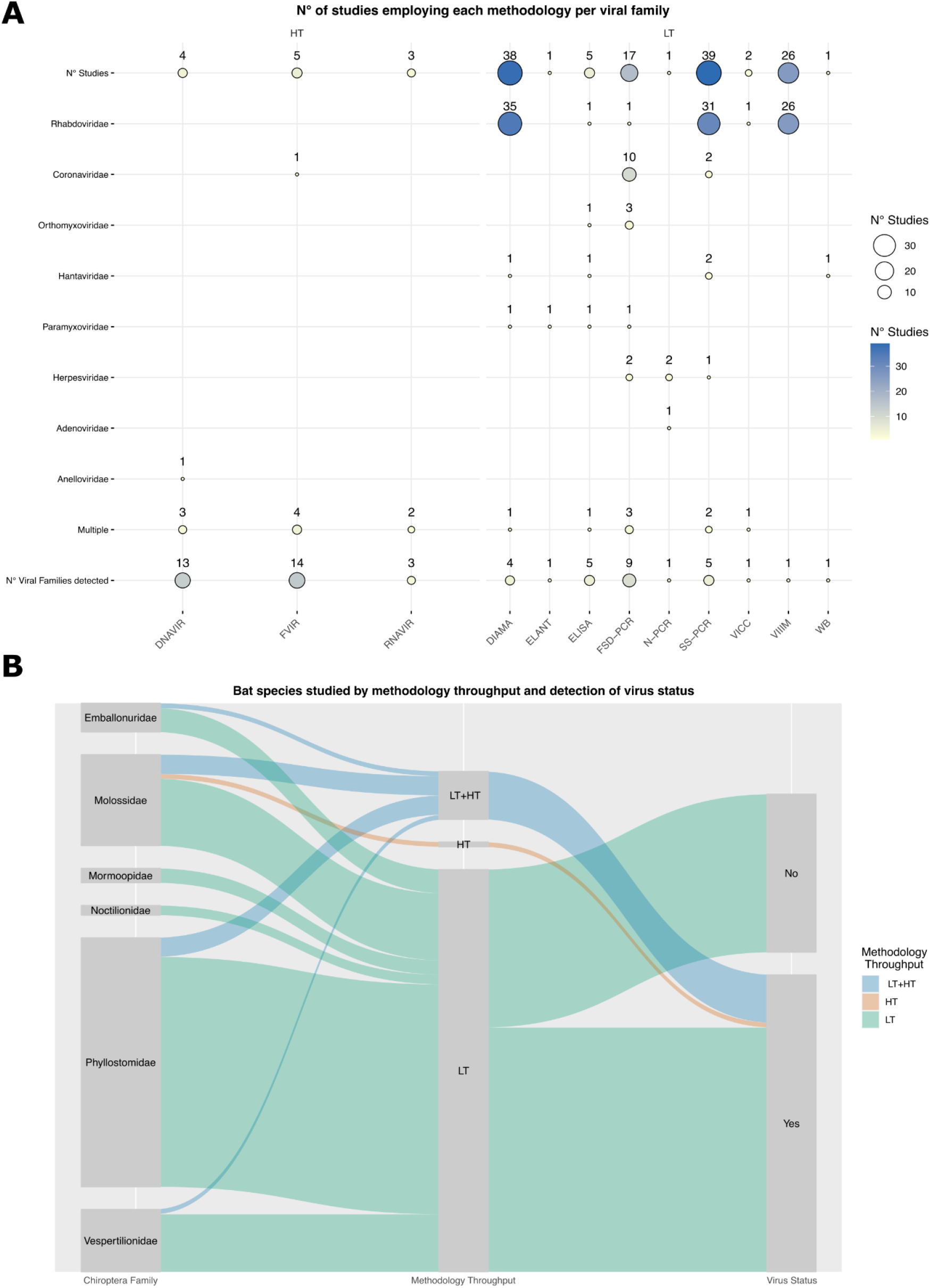
Methodological approaches used to investigate virus and bat species stratified by throughput and infection/exposure status. **A -** Number of studies employing each methodology to investigate specific virus families. **B -** Bat species included in at least one study connected by methodology throughput (LT - low throughput, HT - high throughput, LT+HT low and high throughput) and virus infection/exposure status.

Seventy-five bat species were screened for viruses using LT only, one (*E. bonariensis*) was investigated using a HT method only, and 10 species were investigated using LT and HT (*P, macrotis, D. rotundus, G. soricina, C. perspicillata, A. lituratus, E. patagonicus, M. molossus, M. rufus, T. brasiliensis, E. diminutus*) (**Figure 4B**). Seven of these species (*D. rotundus, G. soricina, C. perspicillata, A. lituratus, M. molossus, M. rufus, T. brasiliensis*) are also amongst the most studied ones (**Figure 1D)**. All 12 bat species that were investigated by at least one HT methodology were found positive (**Figure 4B**), while 33 species investigated using low throughput methodologies were found negative (**Figure 4B**). The negative status of several bat species is likely a result from the methodology chosen for viral characterization, but it may also reflect a composite effect of other sources of bias discussed before, such as sampling, bias towards species and/or tissue sampled, as well as virus/host life history traits such as population/metapopulations infection rate through time.

### Detection and characterization of single or related virus lineages

RABV was extensively characterized via VIIIM and DIAMA and to a lesser extent, these methods have been used for hantaviruses and paramyxoviruses. These methods provide precise information about symptoms induced by RABV infection in mice models and the possibility to isolate abundant viral material. However, it is a laborious methodology requiring euthanasia or sampling sick/dead bats. Therefore, these methodologies are not well suited for more broad virus discovery and virome community characterization.

Only one reviewed study used ELISA to detect Henipa- and filo-like viruses(56). Antibody and antigen detection using ELISA provide only a narrow characterization of viral diversity, restricted to a single or closely related viral lineage. Cross antibody recognition may allow the detection of other viruses, but it only extends to viruses from the same genus and rarely to the same family(57). Moreover, cross-reactivity of serotypes provides little or no information about which other viruses infected or are infecting a given sample, limiting substantially the use of these methods for virus discovery.

Amplicon-based screenings have been applied for diagnosing several viral pathogens mainly because of low cost, high sensitivity and speedy results, allowing high turnover rates. It is also possible to estimate the virus genome copy number and couple it with sequencing of amplified fragments.

One key-constraint of all these methodologies is the need for previous information about viral molecules (proteins, DNA/RNA), which is lacking for the large majority of poorly studied viruses infecting animals(23). Therefore, they are not suited for viral discovery and diversity characterization, but are better suited to obtain detailed information about known viruses.

### Characterization of multiple virus lineages

Degenerated primers targeting conserved regions may allow a broader detection of divergent viruses, however high genetic divergence precludes broad surveys. Nowadays, there are *brute force* methodologies that allow more unbiased detection and characterization of virus genetic material. For instance, high throughput sequencing (HTS) approaches can be applied to characterize the complete genomes of DNA and/or RNA viruses (24). But due to the costs, HTS has been mainly used for a few individuals and species of Brazilian bats. Nevertheless, there is a clear increase in HTS usage for the study of Brazilian bats (**Supplementary File 3A**). This reflects the global shift of the scientific community towards comprehensive characterization of viromes and microbiomes. Moreover, viral enrichment protocols coupled with HTS may provide the most comprehensive and cost-effective available methodological pipeline for unbiased assessment of viral communities, because it substantially reduces the amount of sequenced reads needed to obtain viral genomes (58).

The viral diversity characterization predominantly performed using targeted approaches impose clear limits on our ability to understand the true diversity of viromes in Brazilian bats. Our findings are in close agreement with a recent review describing methodological biases involved in the attempts to broadly characterize coronaviruses worldwide (20).

## Perspectives

The results compiled in this review bring to light the limited understanding of viromes of Brazilian bats. Several biases shapes the greater picture: a) complete lack of studies for almost half of known bat species occurring in the country, b) extensive focus on few zoonotic viruses or virus families, c) sampling design failing to take into account host population size and spatio-temporal dynamics, d) a heterogeneous use of tissue and convenience sampling and e) the deployment of targeted LT methods.

On the other hand, the number of studies using HTS for virome exploration is growing in Brazil and South America. However, they remain focused on a few bat species and small sample sizes due to costs. Therefore, the biases and gaps identified point to our limited basic understanding about the virome of neotropical bats. Uncovering viromes of bats and new zoonotic viruses requires hypothesis-driven experimental design that will reveal not only their core virome, but will also uncover the biotic and abiotic factors modulating infection through time and space. Additionally, it can allow identification of risk factors pertaining to cross-species transmission.

### Call and recommendations for less biased virus diversity and prevalence information and assess new viruses with zoonotic potential

#### Sampling strategy - the importance of bat and virus traits

Bat populations or metapopulations vary widely in size and geographical range, hence appropriate sampling of bat populations should be estimated taking into account these parameters. Infection rate through time is another key factor to take into account while planning sampling design. Prevalence and transmission may vary drastically in space and time, being correlated with the reproductive season and antibody waning in juveniles. Random longitudinal sampling may yield a more comprehensive picture of long-term infections and overall prevalence in a given population. But, this sampling strategy could miss short-lived acute infections in geographically isolated populations. A disease surveillance sampling strategy may be an useful alternative, where the sampling design is planned based on probability of sampling infected individuals (disease prevalence). But it is highly dependent on previous knowledge of disease and the etiological agent which is largely lacking for bats in Brazil.

Comprehensive assessments of virus infections and diversity in bats should use multiple tissue/body fluid collections as much as possible since mechanisms of infection, replication and transmission route vary. However, sampling strategies trying to answer questions about zoonotic potential can also use specific tissues/fluids that are more relevant in the transmission cycle. Coronaviruses, for example, infect respiratory and intestinal tracts in mammals(59), hence oral/nasal fluids as well as excreta are suitable material for learning about infection and shedding dynamics. Moreover, less invasive sampling such as swabbing reduces animal suffering and impact on populations, but may also limit the detection to viruses excreted by specific routes only. There is no ideal sampling strategy for covering all these aspects, but keeping in mind the complexity of the virome and its intricate interactions with the hosts is crucial for designing cost-effective, minimally-invasive studies which can offer comprehensive and relevant results.

#### Sampling strategy - methodologies for the characterization of viromes

Understanding the sensitivity and specificity of different methodologies is of utmost importance for assessing the virome diversity. Different molecular assays can be used for complementary testing, but viral enrichment protocols preceding deep HTS is the most powerful workflow for exploring virus communities. Such methodology will set the stage for *in situ* viral discovery and characterization of core viromes of bats and other animals in the next few years. These HTS technologies allow us to move from questions such as “What can we afford to do virome characterization?” to “What should we do for comprehensive virome characterization?”. Such approaches may for instance allow the sequencing of the virome of every individual sampled, not requiring sample pooling and providing a direct measurement of infection rate and coinfection of several viruses in the same individual. Challenges of HTS should be kept in mind: growing datasets of hundreds of samples are currently amenable only using powerful computational infrastructure and trained researchers(60), substantially limiting its deployment in low resource settings.

A call for application of standardized bat and viral sampling, viral enrichment and bioinformatic protocols is required in order to enable cross-studies comparison.

#### One Health perspective

Direct infection of humans by bat-borne viruses is only known in two specific cases: *Nipah virus* in Bangladesh (61, 62), and RABV in the Americas (34). Otherwise, bat-borne viruses that are able to infect humans also infect a large range of vertebrates including several sylvatic and domestic animals as well as vector species (2). Moreover, there is evidence indicating that these viruses underwent prior adaptation in other mammal hosts before spilling over to humans (59). All these viruses have complex transmission cycles infecting a range of closely and distantly related host species, suggesting that viruses with a large host range are more prone to host switching triggering new outbreaks (10). Therefore, a system-level One Health approach is needed for understanding biotic and abiotic factors shaping transmission dynamics and emergence risk of bat-borne viruses.

## Conclusion

The characterization of Brazilian bats’ viromes is biased toward viruses of zoonotic concern. Moreover, despite the accumulated knowledge regarding these viruses, we have shown that large discrepancies stemming from spatiotemporal sampling bias and the use of techniques unsuitable for comprehensive virome characterisation. Altogether, our review reveals a limited understanding of viromes in bats in Brazil and thus, substantially limiting a proper assessment of the zoonotic potential of most viruses. Ongoing changes of land use in Brazilian biomes (63) represent new opportunities for cross-species virus transmission between humans and wildlife (64, 65). Thus, spillover events from bats and other vertebrates inhabiting Brazilian biomes may lead to outbreaks and epidemics. High throughput techniques for pathogen discovery and surveillance applied to bats from sylvatic-urban interfaces should be prioritized in high risk contact areas.

## Acknowledgment

G.L.W. is supported by the Conselho Nacional de Desenvolvimento Científico e Tecnológico (CNPq) through their productivity research fellowships (303902/2019-1). In addition, G.L.W was also supported by Coodenação de Aperfeiçoamento de Pessoal de Ensino Superior (CAPES) and the Alexander von Humboldt Foundation. E.Ba. is supported by CNPq through a postdoctoral fellowship (152672/2022-2). E.Be. is supported by a CNPq productivity grant. We would like to thank you Lais Ceschini Machado for map rendering assistance.

## Notes

### Competing Interest Statement

The authors have declared no competing interest.

