## Supplementary figures and images for "The virome of bats inhabiting Brazilian biomes: knowledge gaps and biases towards zoonotic viruses"

### Supplementary File 3

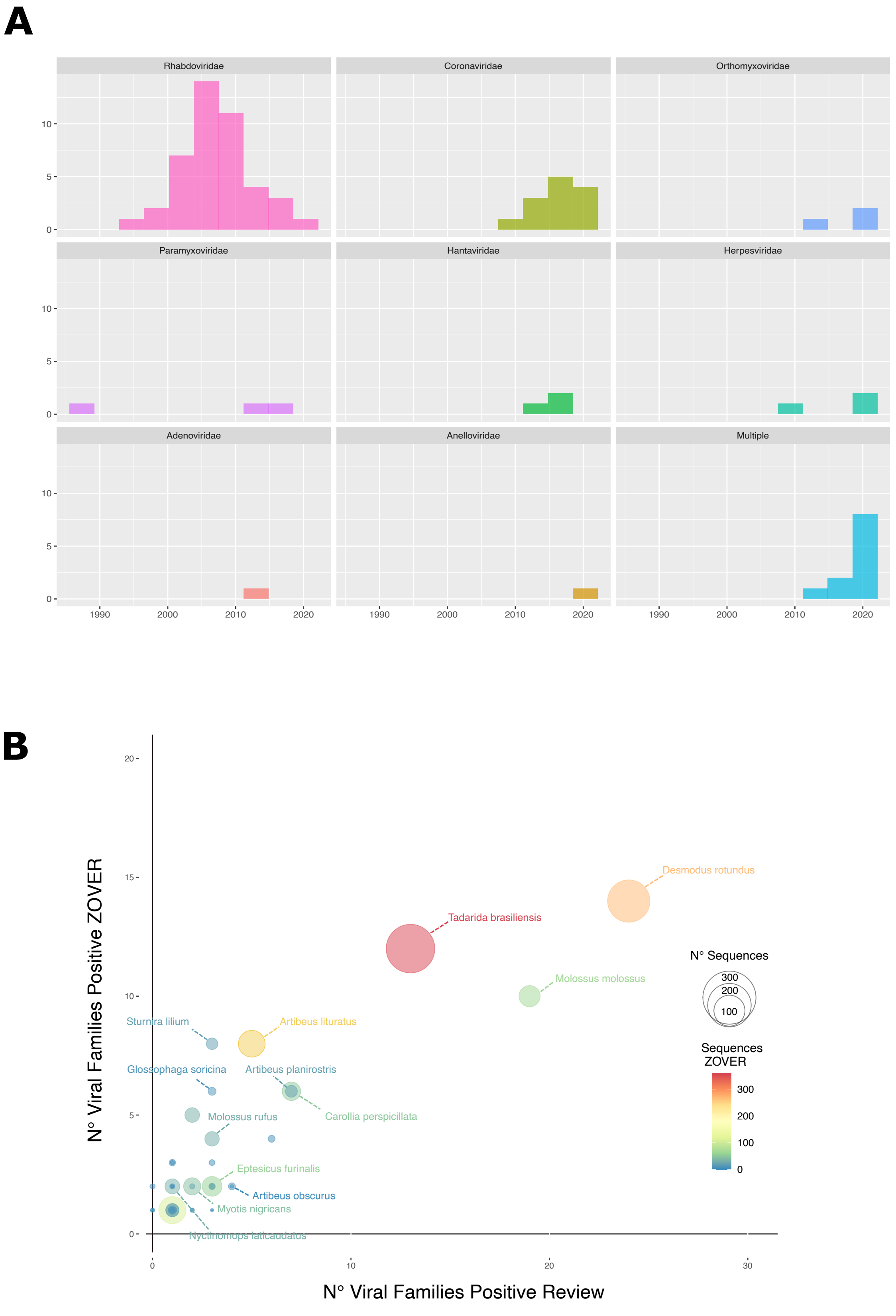

### Supplementary File 5

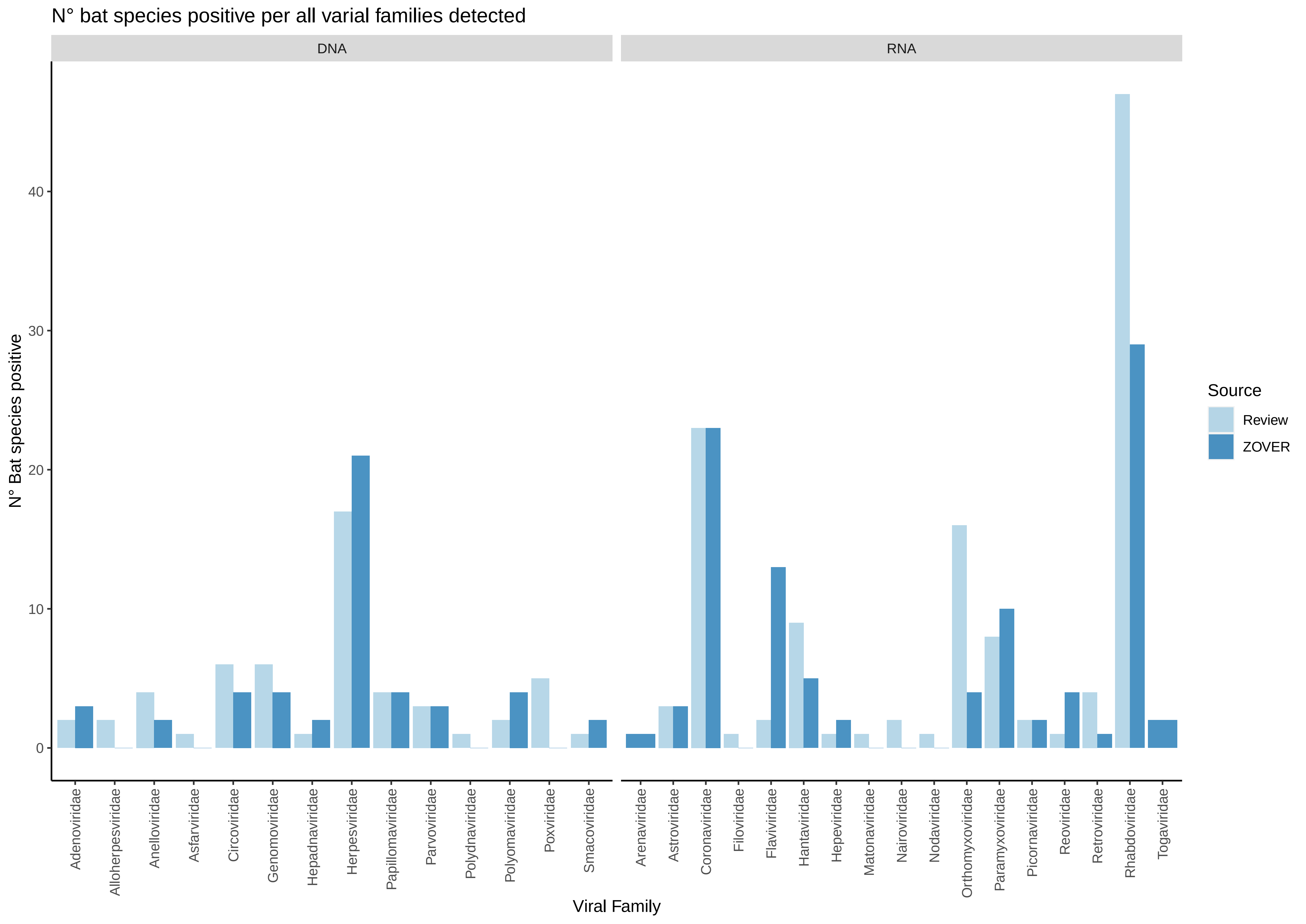
